# HIGD2A is required for assembly of the COX3 module of human mitochondrial complex IV

**DOI:** 10.1101/787721

**Authors:** Daniella H. Hock, Boris Reljic, Ching-Seng Ang, Hayley S. Mountford, Alison G. Compton, Michael T. Ryan, David R. Thorburn, David A. Stroud

## Abstract

Assembly factors play a critical role in the biogenesis of mitochondrial respiratory chain complexes I-IV where they assist in the membrane insertion of subunits, attachment of co-factors, and stabilization of assembly intermediates. The major fraction of complexes I, III and IV are present together in large molecular structures known as respiratory chain supercomplexes. A number of assembly factors have been proposed as required for supercomplex assembly, including the *hypoxia inducible gene 1* domain family member HIGD2A. Using gene-edited human cell lines and extensive steady state, translation and affinity enrichment proteomics techniques we show that loss of HIGD2A leads to defects in the *de novo* biogenesis of mtDNA-encoded COX3, subsequent accumulation of complex IV intermediates and turnover of COX3 partner proteins. Deletion of HIGD2A also leads to defective complex IV activity. The impact of HIGD2A loss on complex IV was not altered by growth under hypoxic conditions, consistent with its role being in basal complex IV assembly. While in the absence of HIGD2A we show that mitochondria do contain an altered supercomplex assembly, we demonstrate it to harbor a crippled complex IV lacking COX3. Our results redefine HIGD2A as a classical assembly factor required for building the COX3 module of complex IV.

## Introduction

The electron transport chain (ETC) of human mitochondria is comprised of four multiprotein complexes (Complex I-IV) embedded in the inner mitochondrial membrane (IMM). In a process known as oxidative phosphorylation the ETC takes electrons from metabolites and uses their energy to build a proton gradient across the IMM, which in turn powers production of ATP through Complex V (F_1_F_o_ ATP Synthase). Complex IV (CIV; CcO; cytochrome *c* oxidase) is the fourth and last enzyme of the ETC and catalyzes the transfer of electrons from cytochrome *c* to molecular oxygen. Complexes I, III and IV have been shown to exist in several higher order structures known as respiratory chain supercomplexes or respirasomes (1–4). These structures have been suggested to reduce production of reactive oxygen species (ROS), improve complex stability and/or distribution throughout the protein dense environment of the IMM, and allow efficient channeling of substrates (3–8).

Complex IV is composed of 14 different subunits of dual genetic origin, the core membrane embedded subunits, COX1, COX2 and COX3 are encoded by mitochondria DNA (mtDNA) genes *MT-CO1, MT-CO2* and *MT-CO3*, that are translated on mitochondrial ribosomes in the matrix, whereas the remaining 11 subunits are encoded by nuclear DNA, translated in the cytosol and imported into the mitochondria where they assemble with mtDNA encoded subunits. Nuclear encoded subunits can therefore be grouped into assembly modules seeded by each of the mtDNA encoded subunits (Fig. 1) (9, 10). During complex IV biogenesis, redox active heme and copper cofactors are incorporated into COX1 and COX2, which together with COX3 form the conserved core of the enzyme. While COX3 does not have a direct role in electron transport, it is thought to play a regulatory role in modulating complex IV activity (11, 12). Homologs of COX3 are present in complex IV throughout the tree of life (13) and mutations within *MT-CO3* lead to complex IV deficiency and mitochondrial encephalopathy or myopathy (MIM 516050; 14, 15) implicating the importance of COX3 in complex IV and mitochondrial function.

**Figure 1.**
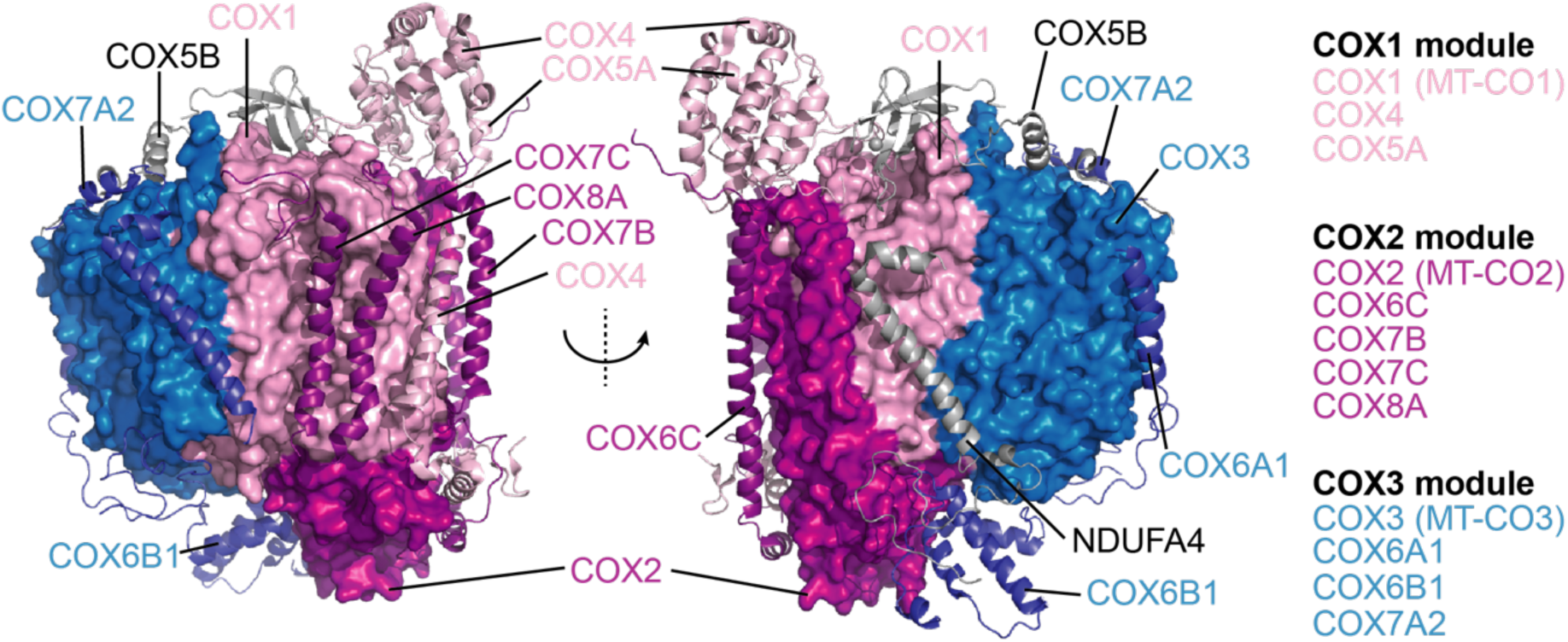
Human complex IV colored by subunit module (PBD 5Z62). Core mtDNA-encoded subunits (COX1 = blue, COX2 = light pink, COX3 = magenta) are rendered as surface representation whereas nuclear DNA-encoded subunits are presented as ribbons.

Biogenesis of complex IV occurs through the coordinated effort of more than 30 known assembly factors with roles including control of mitochondrial RNA stability and translation, membrane insertion of mtDNA encoded subunits, copper and heme biosynthesis and loading and stabilization of subunit modules during assembly (9, 10, 16). While several assembly factors are known to be essential for stabilizing the modules containing COX1 and COX2, to date no assembly factors have been associated with the COX3 module (containing COX3, COX6A, COX6B and COX7A) in humans. Two assembly factors have also been suggested to support complex IV-containing supercomplex assembly, COX7A2L (SCAFI) and *hypoxia inducible gene 1* domain family member HIGD2A (17). COX7A2L was originally shown to be present within the supercomplex but not in the holo form of complex IV (6). A mouse strain harboring a truncated form of the protein lacked some forms of the supercomplexes and exhibited altered mitochondrial respiration, leading the authors to suggest the protein is required for supercomplex stability (6). Several studies have disputed this finding (18–23) and it is now understood that COX7A2L plays a role as a checkpoint for assembly of CIII_2_ into the supercomplex (17), a result that explains the differences in supercomplex distribution observed in mitochondria lacking full length COX7A2L. Yeast (*S. cerevisiae*) homologs of HIGD2A (Rcf1 and Rcf2) belong to the *hypoxia inducible gene 1* domain containing (HIGD) family and were originally suggested to be required for assembly and stability of the yeast CIII_2_/IV supercomplex (24–26). Deletion of Rcf1 leads to impaired assembly of Cox12 and Cox13 (COX6B and COX6A in humans) into complex IV, reduced association with Rcf2, and a reduction in the levels CIII_2_/IV supercomplex (24–26). Recent studies have suggested a regulatory role for Rcf1 and Rcf2 in complex IV activity (27) through association with complex IV intermediates (28). Investigation into the function of mammalian homologs HIGD1A and HIGD2A have been limited to only a few studies. Both *HIGD1A* and *HIGD2A* were originally identified in a screen for genes upregulated during HIF1α dependent hypoxia (29, 30). Following discovery of Rcf1 and Rcf2 in yeast supercomplex assembly, Chen *et al.* (26) showed that knockdown of *Higd1a* in mouse myoblasts had no impact on respiratory chain supercomplexes whereas, knockdown of *Higd2a* led to reduced amounts of CIII_2_/CIV and a higher-order assembly now known to be the respiratory chain megacomplex (CI_2_/III_2_/IV_2_) (2). No impact was observed on the levels of holo complex IV, leading the authors to suggest *Higd2a* is a supercomplex assembly factor, consistent with the studies in yeast (26). In contrast, HIGD1A has recently been suggested to associate non-stoichiometrically with an early assembly intermediate consisting of COX1 and associated subunits (10) and induce structural changes around the heme *a* active center altering complex IV activity during hypoxic stress (31).

We identified HIGD2A as a protein strongly upregulated in lymphoblasts from a patient with ETC complex IV deficiency. Given that the yeast homolog of HIGD2A has been previously associated with either complex IV or respiratory chain supercomplex assembly, we aimed to clarify the function of human HIGD2A as well as its ortholog HIGD1A. Using CRISPR/Cas9 generated knockout cell lines we show that loss of HIGD1A leads to a modest defect in mitochondrial oxygen consumption and stability of subunits present within the COX1 module but does not affect complex assembly. While we find HIGD1A is indeed induced during hypoxia, this results in no apparent changes to complex IV or supercomplex assembly. Conversely, we demonstrate that loss of HIGD2A leads to a strong defect in mitochondrial respiration, complex IV assembly and activity due to impaired stability of newly translated COX3, and subsequent degradation of nuclear encoded subunits present within the COX3 module. We conclude that HIGD2A is a classical assembly factor involved in stabilization of COX3.

## Experimental Procedures

### Tissue culture and generation of cell lines

HEK293T cell lines were cultured in Dulbecco’s modified Eagle’s medium (DMEM High Glucose) supplemented with 10% (v/v) fetal calf serum (FCS; CellSera), penicillin/streptomycin (Gibco) and 50 µg ml^−1^ uridine (Sigma) at 37°C under an atmosphere of 5% CO_2_. Lymphoblasts were cultured under the same conditions except that the medium was RPMI-1640 and the amount of FCS was 20% (v/v). Gene editing was performed in HEK293T (originally purchased from ATCC) and the NDUFV3^KO^NDUFV3^FLAG^ cell line published previously (32). Constructs for CRISPR-Cas9 genome editing were designed using the CHOPCHOP software (33) and oligonucleotides encoding gRNA sequences cloned into the pSpCas9(BB)-2A-GFP (PX458) plasmid (a gift from F. Zhang; Addgene, plasmid 48138; (34)) as previously described (32). The gRNA sequences used were 5’-ATGTCAGTCAAACTTACCAA for HIGD1A and 5’-ACTTACCTATGGGTACCACC for HIGD2A. Constructs were transfected using Lipofectamine LTX (ThermoFisher Scientific) according to manufacturer’s instructions. Single GFP positive cells were sorted into 96 well plates and clonal populations expanded for screening by SDS-PAGE and immunoblotting as described below. To verify the CRISPR/Cas9 induced insertions and deletions (indels) genomic DNA was isolated from candidate clones using the Quick-DNA kit (Zymo Research) according to manufacturer’s specifications. Oligonucleotides provided by the CHOPCHOP software (33) were used for amplification of target regions and cloning into pGEM4Z (35) for M13 primed Sanger sequencing of individual alleles as we have done previously (32). HIGD1A^KO^ contained the following mutations c.[97+2G_97+6Gdel];[97+2G_97+14Gdel];[97_97+3Tdel] predicted to impact splicing. HIGD2A^KO^ contained c.[145del];[131_150del];[133_149del];[138_158del];[143_153+49Gdel] predicted to alter the frame and introduce premature stop codons. NDUFV3^KO^NDUFV3^FLAG^HIGD2A^KO^ contained the mutations c.[127_128insA;136_145del];[142_151del];[141_150del];[144_146del];[144_150del];[143_147 delinsGTGGAACGCCCGGCCTCAGTGAGCGAGCGAGCGCGCAGAGAGGGAGTGGCCA ACTCCATCACTAGGGGTTCCTGCGGCCGCTCCCCAGCATGCCTGCTATTCTCTTCCCA ATCCT] predicted to alter the reading frame and introduce premature stop codons. For hypoxia treatment, cells were cultured inside a hypoxia incubator chamber (STEMCELL Technologies) pre-purged with 1 % O_2_, 5 % CO_2_, 94 % N_2_ for 10 min at 25 l min^-1^. The sealed chamber was incubated at 37°C for 3 or 24 hours as indicated. For pulse-SILAC analysis of mitochondrial translation, cells were incubated in DMEM as above with the overnight addition of 50 ug ml^−1^ chloramphenicol (CAP; Sigma) prior to the start of the experiment. Media was replaced with neat DMEM for SILAC (ThermoFisher Scientific) for 15 minutes following which the media was supplemented with 10% dialyzed FCS (ThermoFisher Scientific), Sodium Pyruvate (Gibco), Glutamax (Gibco), penicillin/streptomycin (Gibco), 3.5 g L^−1^ glucose and 0.1 mg ml^−1^ cycloheximide (Sigma). Following an incubation for 30 minutes at 37°C under an atmosphere of 5% CO_2_, media was further supplemented 1.2 g ml^−1^ L-proline, 285 mg ml^−1^ -^13^C_6_^15^N_2_-L-lysine-HCl and 85 mg L^-1^-^13^C_6_^15^N_2_-L-arginine-HCl (Silantes). Time points were collected at 0, 1, 2 and 4 h post addition of SILAC amino acids.

### Patient sequencing and enzymology assays

DNA was extracted from skin fibroblasts and whole exome sequencing was performed as published (per subject 1 in reference) (36). Electron transport chain enzyme activities in skeletal muscle biopsies (post-660g supernatants) and mitochondrial preparations from Epstein-Barr Virus (EBV) transformed lymphocytes (lymphoblasts) were determined spectrophotometrically as previously described (37). Enzymes assayed were complex I (NADH-coenzyme Q1 oxidoreductase), complex II (succinate-coenzyme Q1 oxidoreductase), complex III (decylbenzylquinol-cytochrome *c* oxidoreductase), complex IV (cytochrome *c* oxidase) plus the mitochondrial marker enzyme citrate synthase.

### Retroviral transduction

To rescue HIGD2A expression in HIGD2A^KO^ a construct encoding HIGD2A with a C-terminal FLAG tag and compatible overhangs for Gibson assembly was commercially synthesized (IDT technologies). The construct was combined with pBABE-puro plasmid (Addgene, 1764) cut with BamHI-HF and HindIII-HF restriction enzymes (NEB) and Gibson assembled using the NEBuilder HiFi DNA Assembly System (NEB) as per manufacturer’s directions to yield pBabe-puro-HIGD2A^FLAG^. The construct was used to generate viral particles in HEK293T cells as previously described (32). Briefly, pBabe-puro-HIGD2A^FLAG^ was co-transfected alongside pUMVC3 and pCMV-VSV-G (Addgene, 8449 and 8454) using Lipofectamine LTX (ThermoFischer Scientific) according to manufacturer’s instructions. Viral supernatant was collected at 48 h post-transfection, filtered with 0.45 μm PVDF membrane (Milipore) and used to infect HIGD2A^KO^ cells in the presence of 8 µg ml^−1^ polybrene. Cells were selected in puromycin at 2 ug ml^−1^ for 10 days. Transduction was verified by SDS-PAGE and immunoblotting.

### SDS-PAGE, BN-PAGE and western blotting

SDS-PAGE was performed using samples solubilized in LDS sample buffer (Invitrogen) with presence of 50 mM DTT and separated on Invitrogen Bolt Bis-Tris protein gels (10% or 4-12%) using the MES running buffer according to manufacturer’s instructions (ThermoFischer Scientific). For assessing the effect of hypoxia treatment, cells were solubilized in RIPA buffer (25mM Tris-HCl pH7.4, 150 mM NaCl, 0.1 % (w/v) SDS, 1% (v/v) Triton X-100) supplemented with 1 mM PMSF and protease inhibitor cocktail (cOmplete™, Merck) on ice. BN-PAGE was performed using mitochondrial protein solubilized in 1% digitonin (w/v) based solubilization buffer as described previously (38, 39). Mitochondria were isolated as previously described (40) with protein concentration determined using the Pierce Protein Assay Kit (ThermoFisher Scientific). Detergent solubilized complexes were separated on Invitrogen NativePAGE Bis-Tris gels (3-12%) as per manufacturer’s instructions. Western blotting of SDS-PAGE gels was undertaken using the Invitrogen iBlot2 Dry Blotting System using PVDF based stacks. The Invitrogen Mini Blot Module transfer system was used to transfer BN-PAGE gels onto PVDF (Millipore) as per manufactures recommendations. Immunoblots were developed using horseradish peroxidase coupled mouse and rabbit secondary antibodies (Cell Signaling Technology) and ECL chemiluminescent substrates (Bio-Rad) and visualized on a ChemiDoc XRS+ gel documentation system (Bio-Rad). Commercial antibodies were obtained for COX1 (Abcam; ab14705), COX2 (Abcam, ab110258), COX4 (Abcam, ab110261), COX6A1 (Proteintech, 11460-1-AP), FLAG (Sigma, F1804), GAPDH (Proteintech, HRP-60004), HIF-1α (Santa Cruz, SC-53546), HIGD1A (Proteintech, 21749-1-AP), HIGD2A (Proteintech, 21415-1-AP), SDHA (Abcam, Ab14715), UQCRC1 (Abcam, AB110252), β-actin (Proteintech, HRP-60008) and used at the recommended dilutions. In-house polyclonal rabbit antibodies for NDUFA9 were previously described (39). In-gel activity assays were performed as described previously (41).

### Oxygen consumption measurements

Mitochondrial oxygen consumption rates were measured in live cells using a Seahorse Bioscience XFe-96 Analyzer (Agilent). Briefly, 25,000 cells were plated per well in culture plates treated with 50 µg/mL poly-D-lysine. For each assay cycle, there were 4 measurements of 2 minutes followed by mixing, 2 minutes wait, and 5 minutes measure. The following inhibitor concentrations were used: 1 µM oligomycin, 1 µM carbonyl cyanide 4-(trifluoromethoxy)phenylhydrazone (FCCP), 0.5 µM rotenone and 0.5 µM antimycin A. Data were normalized using the Pierce Protein Assay Kit (ThermoFisher Scientific) and analyzed using Wave (version 2.6.0, Agilent) and Prism (version 8.0.2, GraphPad) software packages. For substrate driven oxygen consumption, cells were permeabilized in 0.025 % (w/v) digitonin as described (42). For each assay cycle, there were 4 measurements of 2 minutes mix, 2 minutes wait and 5 minutes measure. The following inhibitor concentrations and combinations were used. Port A: TMPD/ascorbate 0.5 mM/2 mM, 1 µM FCCP, ADP 1 mM; Port B: 1 µM oligomycin; Port C: Potassium azide 20 mM. Data were normalized and analyzed as above.

### Acquisition and analysis of steady state mass spectrometry data from patient lymphoblasts

Primary lymphoblasts (two subcultures from the same individual) and controls (from three separate individuals) were pre-normalized based on protein concentration using the Pierce Protein Assay Kit (ThermoFisher Scientific). Pellets were solubilized in 1% (w/v) SDC, 100mM Tris pH 8.1, 40mM chloroacetamide (Sigma) and 10 mM tris(2-carboxyethyl)phosphine hydrochloride (TCEP; BondBreaker, ThermoFisher Scientific) for 5 min at 99°C with 1500 rpm shaking followed by 15 minutes sonication in a waterbath sonicator. Protein digestion was performed with trypsin (ThermoFisher Scientific) at a 1:50 trypsin:protein ratio at 37°C overnight. The supernatant was transferred to stagetips containing 3×14G plugs of 3M™Empore™ SDB-RPS substrate (Sigma) as described previously (32, 43). Ethyl acetate (99% v/v) and 1% (v/v) TFA was added to the tip before centrifugation at 3000 g at room temperature. Stagetips were washed first with the same ethyl acetate containing solution and then subjected to an additional wash with 0.2% (v/v) TFA. Peptides were eluted in 80 % (v/v) acetonitrile (ACN) and 1% (w/v) NH_4_OH, and then acidified to a final concentration of 1% TFA prior to drying in a CentriVap Benchtop Vacuum Concentrator (Labconco) and reconstituted in 0.1 % TFA and 2 % ACN for analysis by Liquid chromatography (LC) - MS/MS on an Orbitrap QExactive Plus (ThermoFisher Scientific) coupled with an Ultimate 3000 HPLC (ThermoFisher Scientific) and NanoESI interface. Peptides were injected onto the trap column (Acclaim C_18_ PepMap nano Trap x 2 cm, 100 μm I.D, 5 μm particle size and 300 Å pore size; ThermoFisher Scientific) at a flow rate of 15 µL/min before switching the trap in-line with the analytical column (Acclaim RSLC C_18_ PepMap Acclaim RSLC nanocolumn 75 μm x 50 cm, PepMap100 C_18_, 3 μm particle size 100 Å pore size; ThermoFisher Scientific). The peptides were eluted using a 250 nL/min non-linear ACN gradient of buffer A (0.1 % FA, 2% ACN) buffer B (0.1 % FA, 80% ACN) increasing from 2.5% to 35.4% followed by a ramp to 99% over 278 minutes. Data were acquired in positive mode using data dependent acquisition 375 – 1800 m/z as a scan range, HCD for MS/MS of the top 12 intense ions with charges ≥ 2. MS1 scans were acquired at 70,000 resolution (at 200 m/z), MS max. injection time of 54 ms, AGC target 3e^6^, NCE at 27%, isolation window of 1.8 Da, MS/MS resolution of 17,500, MS/MS AGC target of 2E^5^. Raw files were processed using the MaxQuant platform (version 1.6.0.16) (44) and searched against UniProt human database (Aug 2017) and a list of common contaminants using default settings for a label free quantification (LFQ) experiment with “LFQ” and “Match between runs” enabled. For this search Trypsin/P cleavage specificity (cleaves after lysine or arginine, even when proline is present) was used with a maximum of 2 missed cleavages. Oxidation of methionine and N-terminal acetylation were specified as variable modifications. Carbamidomethylation of cysteine was set as a fixed modification. A search tolerance of 4.5 ppm was used for MS1 and 20 ppm for MS2 matching. False discovery rates (FDR) were determined through the target-decoy approach set to 1% for both peptides and proteins. The proteinGroups.txt output from the search was processed in Perseus (version 1.6.2.2) (45). Briefly, Log_2_-transformed LFQ Intensities were grouped into patient or control groups consisting of three controls and two replicates of the patient lymphoblasts. Identifications filtered to include only those that were quantified in at least two samples from each group. Annotations for proteins present in the Mitocarta2.0 dataset (46) were added through matching by gene name. A two-sided t-test with significance determined by permutation-based FDR statistics (FDR 5%, S=0.1) was performed with results expressed as a volcano plot.

### Acquisition and analysis of steady state mass spectrometry data from HEK293T cells

Crude mitochondrial protein from HEK293T, HIGD2A^KO^ and HIGD1A^KO^ cells and whole cell material from hypoxia treated HEK293T cells (derived from two independent subcultures of each) were pre-normalized using the Pierce Protein Assay Kit (ThermoFisher Scientific). Protein pellets were solubilized and digested into tryptic peptides for mass spectrometry using the iST-NHS kit (PreOmics GmbH) as per manufacturer instructions. Peptides were labelled with 6plex Tandem Mass Tags (TMT) (ThermoFisher Scientific) in 8:1 label:protein ratio as per manufacturer instructions. Pooled samples were fractionated using the Pierce High pH Reversed-Phase Peptide Fractionation Kit (ThermoFisher Scientific) as per manufacturer’s instructions. Individual fractions were dried using a CentriVap Benchtop Vacuum Concentrator (Labconco) and reconstituted in 2 % (v/v) acetonitrile (ACN) and 0.1 % (v/v) trifluoroacetic acid (TFA). Liquid chromatography (LC) coupled MS/MS was carried out on an Orbitrap Lumos mass spectrometer (ThermoFisher Scientific) with a nanoESI interface in conjunction with an Ultimate 3000 RSLC nanoHPLC (Dionex Ultimate 3000). The LC system was equipped with an Acclaim Pepmap nano-trap column (Dionex-C18, 100 Å, 75 µm x 2 cm) and an Acclaim Pepmap RSLC analytical column (Dionex-C18, 100 Å, 75 µm x 50 cm). The tryptic peptides were injected to the trap column at an isocratic flow of 5 µL/min of 2% (v/v) CH_3_CN containing 0.1% (v/v) formic acid for 5 min applied before the trap column was switched in-line with the analytical column. The eluents were 5% DMSO in 0.1% v/v formic acid (solvent A) and 5% DMSO in 100% v/v CH_3_CN and 0.1% v/v formic acid (solvent B). The flow gradient was (i) 0-6min at 3% B, (ii) 6-95 min, 3-22% B (iii) 95-105min 22-40% B (iv) 105-110min, 40-80% B (v) 110-115min, 80-80% B (vi) 115-117min, 80-3% and equilibrated at 3% B for 10 minutes before the next sample injection. The Synchronous Precursor Selection (SPS)–MS3 based TMT method was used. In brief, a full MS1 spectra was acquired in positive mode at 120 000 resolution scanning from 380-1500 m/z. AGC target was at 4e^5^ and maximum injection time of 50ms. Precursors for MS2/MS3 analysis were selected based on a Top 3 second method. MS2 analysis consists of collision induced dissociation (CID) with isolation window of 0.7 in the quadrupole. CID was carried out with normalized collision energy of 35 and activation time of 10 ms with detection in the ion trap. Following acquisition of each MS2 spectrum, multiple MS2 fragment ions were captured in the MS3 precursor population using isolation waveforms frequency notches. The MS3 precursors were fragmented by high energy collision-induced dissociation (HCD) and analyzed using the Orbitrap with scan range of 100-500 m/z, 60 000 resolution, normalized collision energy of 60%, AGC target of 1e^5^ and maximum injection time of 120ms. Raw files were processed using the MaxQuant platform (version 1.6.3.3) (44) and searched against UniProt human database (January 2019) using default settings for a TMT 6plex experiment with the following modifications: deamination (NQ), oxidation of methionine and N-terminal acetylation were specified as variable modifications. Trypsin/P cleavage specificity (cleaves after lysine or arginine, even when proline is present) was used with a maximum of 2 missed cleavages. Carbamidomethylation of cysteine was set as a fixed modification. A search tolerance of 4.5 ppm was used for MS1 and 20 ppm for MS2 matching. False discovery rates (FDR) were determined through the target-decoy approach set to 1% for both peptides and proteins. The proteinGroups.txt output from the search was processed in Perseus (version 1.6.2.2) (45). Briefly, Log_2_-transformed TMT reporter intensity corrected values were grouped into wild-type, HIGD2A and HIGD1A groups consisting of two replicates each. Identifications filtered to include 100% valid values across all samples. Annotations for proteins present in the Mitocarta2.0 dataset (46) were added through matching by gene name and rows filtered to include only mitochondrial entries. A two-sided t-test with significance determined by permutation-based FDR statistics (FDR 5%, S=1) was performed with results expressed as a volcano plot. Ratios for individual subunits relative to controls were mapped onto the Cryo-EM PDB maps cited in the figure legends using a custom python script described previously (32).

### Acquisition and analysis of pulse-SILAC mass spectrometry data

Mitochondrial protein pellets (from three independent subcultures) were solubilized in 1% (w/v) SDC, 100mM Tris pH 8.1, 40mM chloroacetamide (Sigma) and 10 mM tris(2-carboxyethyl)phosphine hydrochloride (TCEP; BondBreaker, ThermoFisher Scientific) for 5 min at 99°C with 1500 rpm shaking followed by 15 minutes sonication in a waterbath sonicator. Protein digestion was performed with trypsin (ThermoFisher Scientific) at a 1:50 trypsin:protein ratio at 37°C overnight. The supernatant was transferred to stagetips containing 3×14G plugs of 3M™Empore™ SDB-RPS substrate (Sigma) as described previously (32, 43) and above. Analysis of reconstituted peptides was performed by LC MS/MS on a QExactive Plus Orbitrap mass spectrometer (ThermoFisher Scientific) in conjunction with an Ultimate 3000 RSLC nanoHPLC (Dionex Ultimate 3000). The LC system was as described above (*Acquisition and analysis of steady state mass spectrometry data from HEK293T cells*). For this experiment the mass spectrometer was operated in the data-dependent mode with a targeted inclusion list containing predicted peptides from the 13 mitochondrial DNA-encoded proteins. The inclusion list consists of mass/charge (m/z) and charge (z) of tryptic peptides (endogenous and SILAC labelled) predicted from *in-silico* digest of target proteins using the Skyline software (47). In addition, the inclusion list also contained peptides that have been previously observed in public data depositories through the PeptideAtlas site (48) and the present study. Full MS1 spectra were acquired in positive mode, 140,000 resolution, AGC target of 3e^6^ and maximum IT time of 50 ms. A loop count of 10 on the most intense targeted peptide were isolated for MS/MS. The isolation window was set at 1.2 m/z and precursors fragmented using stepped normalized collision energy of 28, 30 and 32. Resolution was at 35,000 resolution, AGC target at 2e^5^ and maximum IT time of 150 ms. Dynamic exclusion was set to be 30 seconds. Raw files were processed using the MaxQuant platform (version 1.6.3.3) (44) and searched against UniProt human database May 2018 using default settings for a SILAC experiment. For this search Trypsin/P cleavage specificity (cleaves after lysine or arginine, even when proline is present) was used with a maximum of 2 missed cleavages. Oxidation of methionine and N-terminal acetylation were specified as variable modifications. Carbamidomethylation of cysteine was set as a fixed modification. A search tolerance of 4.5 ppm was used for MS1 and 20 ppm for MS2 matching. False discovery rates (FDR) were determined through the target-decoy approach set to 1% for both peptides and proteins. The “Requant” option was set to “on”. The proteinGroups.txt output from the search was processed in Perseus (version 1.6.2.2) (45). Briefly, heavy intensity values for the detected mtDNA encoded proteins were Log_2_ transformed and values were normalized to the maximum value detected in the control 4 h condition. The means from three experiments were plotted over time using Prism (version 8.1.2, GraphPad) along with the standard deviation from the mean of replicates.

### Acquisition and analysis of affinity enrichment mass spectrometry data

Affinity enrichment experiments were performed on complemented HIGD2A^FLAG^ cells. In brief, 1 mg of whole-cell protein pellets from HEK293T and HIGD2A^FLAG^ (derived from three subcultures of each) were harvested in triplicate and solubilized in 20mM Tris-Cl pH 7.4, 50mM NaCl, 10% (v/v) glycerol, 0.1mM EDTA, 1% (w/v) digitonin and 125 units of benzonase (Merck). Pierce™ Spin Columns (ThermoFisher Scientific) were loaded with 40 μl anti-FLAG M2 affinity gel (Sigma) and equilibrated with wash buffer (20mM Tris-Cl pH 7.4, 60mM NaCl, 10% v/v glycerol, 0.5mM EDTA, 0.1% w/v digitonin). Clarified (20,000 x g, 5 mins at 4°C) lysate was added to spin columns containing FLAG beads and incubated at 4°C for 2 h in rotating mixer. Columns were attached to a vacuum manifold and washed 30 times with 500 μl of wash buffer. For elution, 100 μl of wash buffer containing 100 μg ml^−1^ of FLAG peptide; Sigma) for 30 minutes at 4°C prior to low speed centrifuge and collection into a microcentrifuge tube. Another 100 μl of wash buffer (not containing FLAG peptide) was added and both elutions were pooled into the same tube. Eluted proteins were precipitated with 5x volume of ice-cold acetone overnight at −20°C. On the next day, samples were centrifuges at max speed for 10 minutes at room temperature and the precipitated protein pellet was solubilized with 8 M urea in 50 mM ammonium bicarbonate (ABC) followed by sonication for 15 minutes in a sonicator waterbath. To reduce and alkylate proteins tris(2-carboxyethyl)phosphine hydrochloride (TCEP; Bondbreaker, ThermoFisher Scientific) and chloroacetamide (Sigma Aldrich) were added to a final concentration of 10mM and 50mM respectively and incubated at 37°C for 30 minutes while shaking. Samples were diluted to 2M urea using 50 mM ABC prior to digestion with 1 μg of trypsin (ThermoFisher Scientific) at 37°C overnight. Peptides were acidified to 1% TFA and desalted using stagetips (43) containing 2x 14G plugs of 3M™ Empore™ SDB-XC Extraction Disks (Sigma) which were pre-activated with 100% acetonitrile (ACN) and equilibrated with 0.1% TFA, 2% ACN prior to sample loading. Column was washed with 0.1% TFA, 2% ACN and samples were eluted with 80% ACN, 0.1% TFA. All centrifugation steps were performed at 1,800 g. Elutions were dried using CentriVap concentrator (Labconco) and samples were reconstituted in 0.1% TFA, 2% ACN. LC-MS/MS was carried out on a LTQ Orbitrap Elite (Thermo Scientific) in conjunction with an Ultimate 3000 RSLC nanoHPLC (Dionex Ultimate 3000). The LC system was as described above (*Acquisition and analysis of steady state mass spectrometry data from HEK293T cells*). The LTQ Orbitrap Elite was operated in the data-dependent mode with nanoESI spray voltage of 1.8kV, capillary temperature of 250°C and S-lens RF value of 55%. All spectra were acquired in positive mode with full scan MS spectra from m/z 300-1650 in the FT mode at 120,000 resolution. Automated gain control was set to a target value of 1.0e^6^. Lock mass of 445.120025 was used. The top 20 most intense precursors were subjected to rapid collision induced dissociation (rCID) with normalized collision energy of 30 and activation q of 0.25. Dynamic exclusion with of 30 seconds was applied for repeated precursors. Raw files were analyzed using MaxQuant platform (44) (version 1.6.5.0) and searched against UniProt human database January 2019 using default LFQ search parameters with the following modifications: LFQ min. ratio count and label min. ratio count = 1. For this search Trypsin/P cleavage specificity (cleaves after lysine or arginine, even when proline is present) was used with a maximum of 2 missed cleavages. Oxidation of methionine and N-terminal acetylation were specified as variable modifications. Carbamidomethylation of cysteine was set as a fixed modification. A search tolerance of 4.5 ppm was used for MS1 and 20 ppm for MS2 matching. False discovery rates (FDR) were determined through the target-decoy approach set to 1% for both peptides and proteins. Data analysis was performed using Perseus framework (45) (version 1.6.2.2) Briefly, LFQ intensities imported from the proteinGroups.txt output were Log_2_ transformed. Only proteins quantified across all three IP samples (with no restrictions on control identifications) were included and the missing control values imputed at the limit of detection based on normal distribution. Experimental groups were assigned to each set of triplicates and a two-sided t-test performed with significance determined by permutation-based FDR statistics (FDR 5%, S=1) to exclude all identifications enriched in control samples.

### Experimental Design and Statistical Rationale

For label-free and TMT-labelled analyses of whole cells or mitochondria, the statistical approaches used to analyze the data were consistent with published analyses from ours (32, 49) and other labs employing similar instrumentation and methods. The log2 ratio values were normally distributed. The fold change threshold we used for significance was determined through a modified two-sided t-test based on permutation-based FDR statistics (REF) between the two groups, with an FDR of 5% and the s0 value determined by the main distribution of quantified proteins. AEMS experiments were performed in triplicate using LFQ and compared to control cells as we have done previously (32, 49). Imputation was applied only to controls and random values drawn from a distribution equivalent to the limit of detection in each experiment. The log2 intensity values were normally distributed. Significantly enriched proteins were determined through a modified two-sided t-test based on permutation-based FDR statistics (45) between the two groups, with an FDR of 5% and the s0 value set to exclude enriched proteins specific to the control group. For the pulse-chase SILAC experiment rows for mtDNA encoded proteins were isolated and log2 heavy proteinGroup intensities scaled to the maximum intensity observed for the proteinGroup in control cells.

## Results

### HIGD2A is upregulated in patient lymphoblasts with an isolated complex IV deficiency

We identified a patient presenting with intrauterine growth retardation, developmental delay, cortical blindness, possible deafness, lactic acidosis and cardiomyopathy. The patient died at 16 months of age. Enzymology revealed an apparently isolated complex IV defect in muscle and lymphoblasts (Table 1). Whole exome sequencing revealed two novel compound heterozygous missense mutations (c.1874G>A, p.Arg625His; c.665C>T p.Thr222Ile) in *AARS2,* which encodes for mitochondrial alanine-tRNA synthetase (MIM 612035). Both mutations are classified as class 4 or ‘likely pathogenic’ (PM2, PP3, PS3) using the ACMG standards and guidelines for the interpretation of sequence variants (50). It is not uncommon to find an apparently isolated ETC enzyme defect in genetic defects that would be expected to affect multiple ETC enzymes (51). Given that *AARS2* lymphoblasts harbor an isolated complex IV defect, we utilized them to search for novel proteins involved in assembly of complex IV. We performed quantitative mass spectrometry on the patient lymphoblasts and on control lymphoblasts from three unaffected individuals, as we have done previously for patient fibroblasts (36). We quantified the levels of 3,901 cellular proteins detected in *AARS2* lymphoblasts and in at least two out of three control cell lines (Supplemental Table S1), revealing significant reduction in the levels of complex IV subunits but not subunits of complexes I, III or V (Fig. 2) consistent with enzymology data. HIGD2A was the most significantly upregulated protein in *AARS2* lymphoblasts relative to control cell lines.

**Figure 2.**
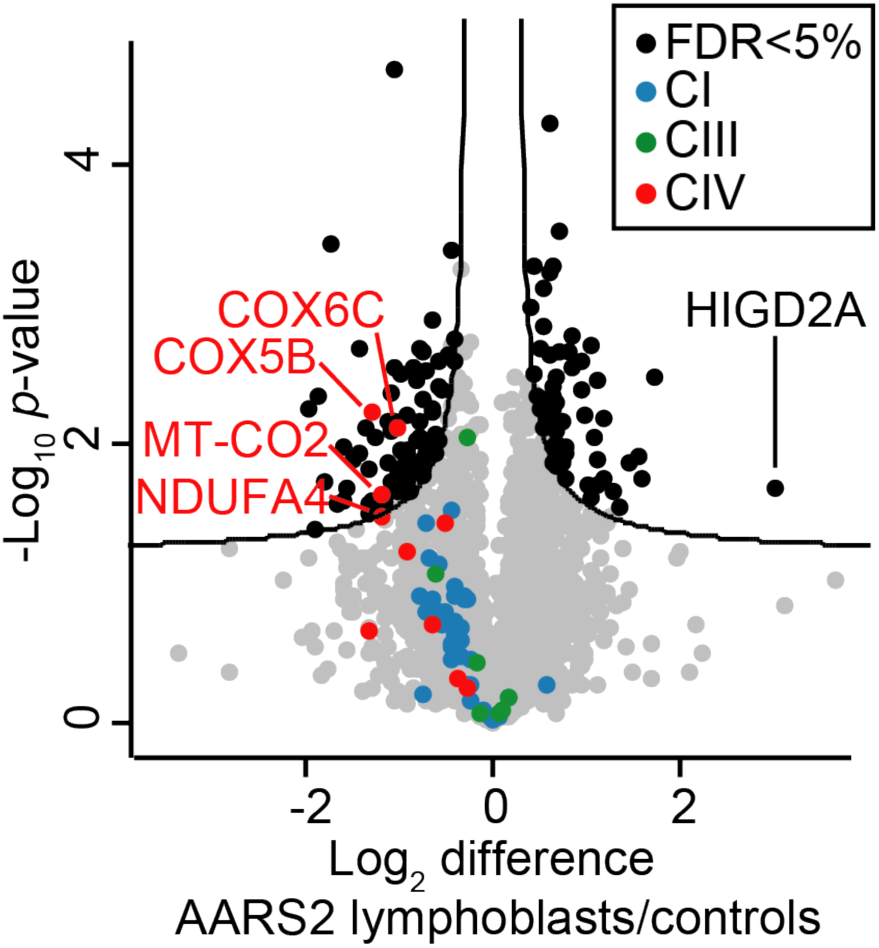
HIGD2A is upregulated in lymphoblasts with a specific defect in the translation of mtDNA-encoded complex IV subunits. Volcano plot comparing protein abundances for proteins quantified in *AARS2* lymphoblasts relative to control lymphoblasts. Subunits of complexes I, III and IV are colored as indicated. The curved line indicates significantly altered protein abundance determined through a false discovery rate-based approach (5% FDR, s0=0.1). n=2 subcultures for the patient cells and n = 3 independent controls.

**Table 1.**
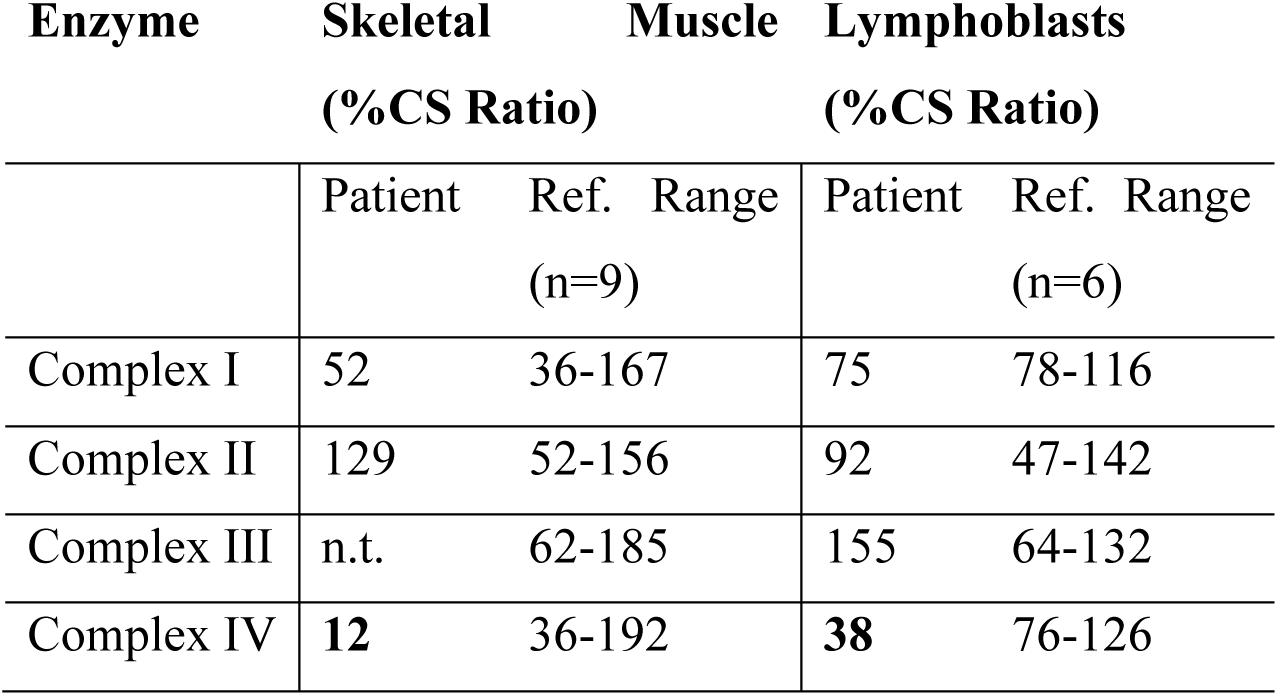
Primary human lymphoblasts with mutated *AARS2* have an isolated complex IV defect. Electron transport chain enzyme activities in skeletal muscle homogenate and EBV-Lymphoblast mitochondria are expressed as %CS ratio, which represents % of the normal control mean value when expressed relative to the mitochondrial marker enzyme Citrate Synthase. Bold characters indicate clinically significant abnormal values. n.t.; not tested

### Loss of HIGD2A leads to defective assembly of complex IV

To clarify the function of HIGD2A we used CRISPR/Cas9 to generate human embryonic kidney (HEK293T) cell lines lacking its expression (Fig. S1). We also generated cell lines lacking the HIGD2A ortholog HIGD1A. Loss of HIGD1A (HIGD1A^KO^) led to no changes in the steady state level of complex IV subunits (Fig. S1A), whereas loss of HIGD2A (HIGD2A^KO^) led to a strong reduction in the levels of COX2 (Fig. S1B), the core subunit of the COX2 module, and the total loss of COX6A, a nuclear encoded subunit present within the COX3 module. To assess the assembly of complex IV and supercomplexes we performed Blue Native (BN)-PAGE. When using the weak non-ionic detergent digitonin, BN-PAGE is capable of resolving stalled complex IV assembly intermediates, holo complex IV and multiple high-order supercomplex structures (10, 17, 20, 52, 53). As can be seen in Fig. 3A, HIGD2A^KO^ mitochondria contain reduced levels of the primary supercomplex assembly consisting of CI/III_2_/IV and accumulation of the CI/III_2_ assembly lacking complex IV (compare lanes 1,2, and 2,5). Additionally, HIGD2A^KO^ mitochondria accumulate both the S2 intermediate containing COX1 and COX4 (Fig. 3A compare lanes 7,8 and 10,11) and S3 intermediate containing COX2, consistent with defective assembly of complex IV (9, 16, 53). Strikingly, no signal was observed for COX3 module subunit COX6A1, either in accumulated intermediates or mature complex IV assemblies (Fig. 3A compare lanes 16,17). In contrast to HIGD2A^KO^, mitochondria lacking HIGD1A contained normal levels and distribution of supercomplex assemblies and accumulated complex IV intermediates were not detected (Fig. 3A).

**Figure 3.**
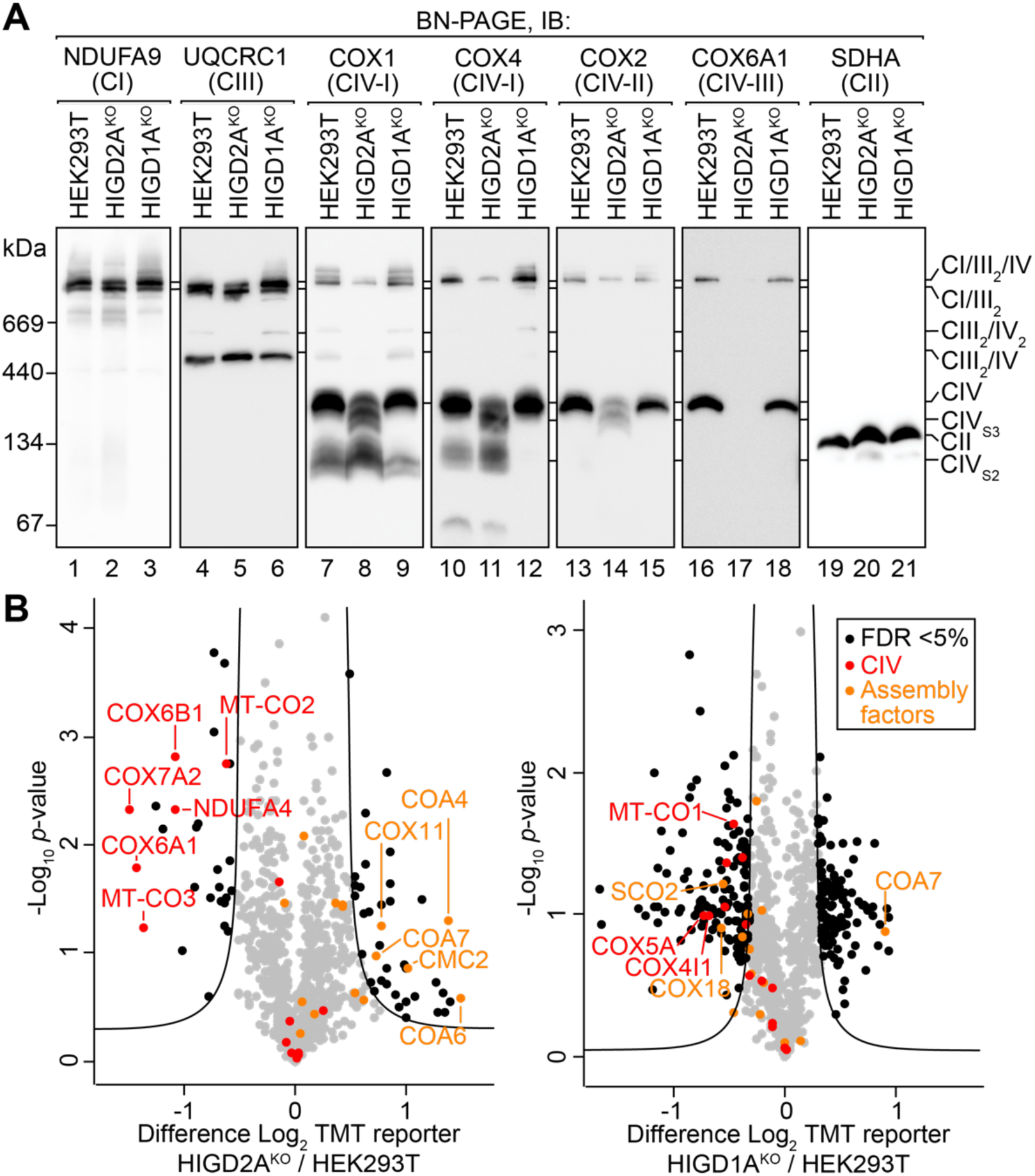
Loss of HIGD2A leads to defects in assembly of complex IV. (A) Mitochondria isolated from the indicated cell lines were solubilized in 1% digitonin, analyzed by BN-PAGE and immunoblotted (IB) using antibodies specific for the indicated antibodies. CIV_S2_, complex IV stage II intermediate; CIV_S3_, complex IV stage 3 intermediate. (B) Volcano plots depicting relative levels of mitochondrial proteins in HIGD2A^KO^ and HIGD1A^KO^ cell lines compared to HEK293T cells. Black dots and the curved line represent significantly altered protein abundance determined through a false discovery rate-based approach (5% FDR, s0=1). n=2 independent subcultures.

### HIGD2A stabilizes the COX3 module of complex IV

To confirm the steady levels of complex IV subunits in HIGD2A^KO^ mitochondria we performed quantitative mass-spectrometry. Mitochondrial proteins from HIGD2A^KO^, HIGD1A^KO^ and control cells were digested, and tryptic peptides labeled with tandem mass tags (TMT) prior to pooled analysis and synchronous precursor selection (SPS)–MS3 based acquisition. We quantified the levels of 892 proteins present within the Mitocarta2.0 (46) inventory of mitochondrial proteins across all samples (Supplementary Table S2). As can be seen in Fig. 3B (left panel) HIGD2A^KO^ mitochondria contained a significant reduction in the levels of complex IV subunits COX3 (MT-CO3), COX6A1, COX6B1, COX7A2, as well as COX2 (MT-CO2) and NDUFA4, a subunit originally ascribed to complex I but now known to be present within complex IV (52). Other complex IV subunits were detected but found not to be significantly altered relative to control cells. HIGD2A^KO^ mitochondria also contained significantly increased levels of complex IV assembly factors COA4, COA6, COA7, CMC2 and COX11 (9, 16). In contrast our, analysis of HIGD1A^KO^ mitochondria revealed a significant reduction in the levels of a different cohort of complex IV subunits; COX1 (MT-CO1), COX4 and COX5A, as well as assembly factors SCO2 and COX18 (Fig. 3B, right panel). COA7 was also found in HIGD1A^KO^ mitochondria at increased abundance relative to controls. Notably, the fold change in abundance of complex IV subunits was less in HIGD1A^KO^ (∼1-1.5 fold change) than for HIGD2A^KO^ (∼2-3 fold change) consistent with the strong defect in assembly we observed for the latter (Fig. 3A).

We used topographical heatmaps (32) to interpret our mass-spectrometry data in the context of the intact 14-subunit human holo complex IV (54) and CI/III_2_/IV respiratory chain supercomplex (2) structures recently determined by single particle cryo-electron microscopy. Both structures were determined using complexes isolated from the same HEK293 derived human cell line used in our study. Unsupervised hierarchical clustering of knockout to control protein ratios from both cell lines revealed several clusters of co-dependent subunits from complexes I, III and IV (Fig. 4A). The most strongly clustered cohort of proteins in HIGD2A^KO^ belonged to the COX3 module (Figs. 4A, B). We found NDUFA4 to be strongly correlated with these proteins, suggestive of it being a subunit of the COX3 module. Mapping of our data to the supercomplex (Fig. 4C) revealed the affected complex I and III subunits to be predominantly located at contact sites with complex IV. In contrast, loss of HIGD1A led to the comparatively weak clustering of COX1 and COX2 module subunits (Figs. 4A, D). Taken together, our data suggest that loss of HIGD2A strongly affects the COX3 module of complex IV, leads to accumulation of a crippled complex IV and its defective association with complexes I and III in the respiratory chain supercomplex.

**Figure 4.**
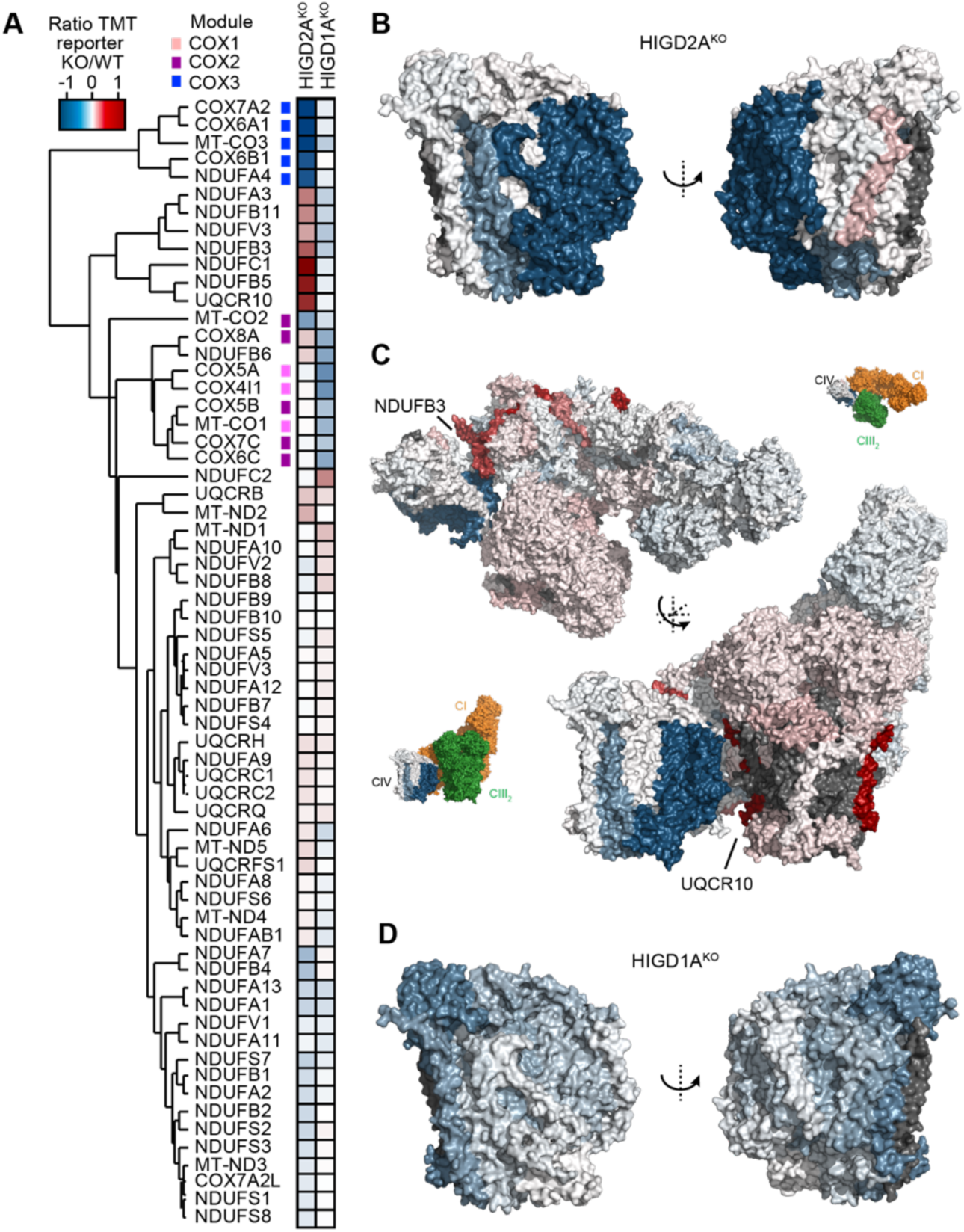
Loss of HIGD2A affects the COX3 module of complex IV. (A) Two-dimensional heatmap showing hierarchically clustered protein ratios for subunits of complex I, III and IV. Modules of complex IV are indicated with the colored squares. (B) Topographical heatmap showing protein ratios for subunits of complex IV in HIGD2A^KO^ cells (PBD 5Z62) (C) Topographical heatmap showing protein ratios for subunits of the human supercomplex in HIGD2A^KO^ cells (PBD 5XTH). The insets show the orientation of the complex with complexes I and III colored in orange and green respectively. (D) Topographical heatmap showing protein ratios for subunits of complex IV in HIGD1A^KO^ cells (PBD 5Z62).

### Loss of HIGD2A leads to impaired complex IV activity

HIGD2A^KO^ cells die in media where glucose has been replaced with galactose (data not shown), indicative of severely impaired mitochondrial respiration (32, 53). To investigate the impact of HIGD2A and HIGD1A on mitochondrial energy generation we measured the oxygen consumption rates in our HIGD2A^KO^ and HIGD1A^KO^ cell lines. Cells lacking HIGD2A were significantly impaired in basal and maximal respiration (Fig. 5A). In contrast, HIGD1A^KO^ had impaired basal respiration but similar maximal respiration compared to control cells leading to an increase in spare respiratory capacity. To investigate the source of the respiration defects we performed an in-gel complex IV activity assay (41). As can be seen in Fig. 5B, the remnant complex IV found within HIGD2A^KO^ mitochondria showed only background levels of activity. While activity was detected in the HIGD2A^KO^ CI/III_2_/IV supercomplex it was reduced relative to control cells. In contrast, complex IV activity in holo complex IV as well as the supercomplex was unaltered in HIGD1A^KO^ mitochondria. Finally, to confirm the impact of HIGD2A loss on complex IV activity *in vivo* we measured complex IV-linked respiration (42). Briefly, the electron donor *N*,*N*,*N*′,*N*′-Tetramethyl-*p*-phenylenediamine (TMPD) is provided in the presence of ascorbate to semi-permeabilized cells where it donates electrons to cytochrome *c* thereby bypassing complex III (Fig. 5C). When TMPD was added to control HEK293T cells in the presence of ADP and uncoupler FCCP, we observed maximal mitochondrial respiration that could be partially dissipated with oligomycin. Inhibition of complex IV activity with azide reveals complex IV dependent oxygen consumption. In contrast to control cells, HIGD2A^KO^ cells had significantly impaired complex IV dependent respiration (Fig. 5C). Taken together, we conclude that loss of HIGD2A leads to a specific defect in complex IV enzymatic activity leading to impaired mitochondrial respiration.

**Figure 5.**
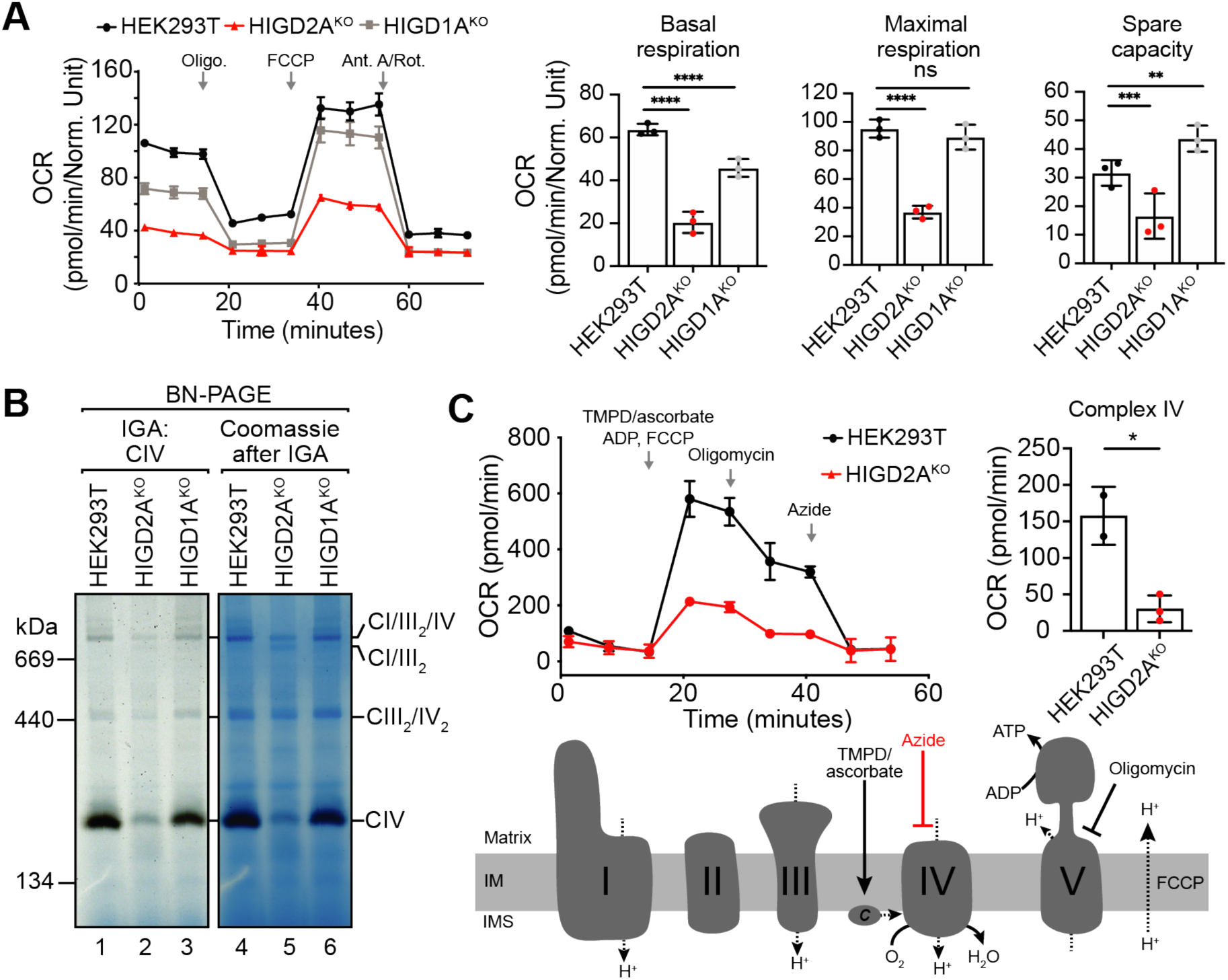
Biochemical characterization of mitochondrial respiration and complex IV activity in HIGD2A^KO^ cells. (A) Oxygen consumption rates (OCR) of HIGD2A^KO^, HIGD1A^KO^ and control cell lines. Data reported as mean ± S.D. n=5. **, p<0.01; ***, p<0.001; ****, p<0.0001. (B) Mitochondrial complexes solubilized in 1% digitonin were separated by BN-PAGE and the activity of complex IV determined through an in-gel assay. The gel was subsequently stained with coomassie (right panel). (C) Flux analysis of complex IV activity in HIGD2A^KO^. OCR was determined in semi-permeabilized cells with the addition of electron donor *N*,*N*,*N*′,*N*′-Tetramethyl-*p*-phenylenediamine (TMPD) and ascorbate that donate electrons to cytochrome *c* bypassing complex III (see schematic). The presence of ADP and FCCP permit maximal respiration. Complex V is inhibited by the addition of oligomycin and complex IV through azide revealing complex IV activity (top right). Data reported as mean ± S.D. n=2 and 3, *, p<0.05.

### HIGD2A acts on complex IV assembly independent of oxygen levels

Mammalian HIGD1A and HIGD2A were originally discovered as proteins induced following hypoxic stress (29, 30), an insult that is known to ultimately lead to turnover of OXPHOS subunits, a reduction in oxidative capacity and an increase in glycolysis. As a way of compensation to these changes in conditions, during the initial stages of hypoxia HIGD1A has been proposed to bind complex IV and promote its activity (31). To address whether the presence or absence of HIGD2A impacts complex IV under hypoxia we cultured HEK293T cells under an atmosphere of 1% oxygen for 0, 3 or 24 hours. Growth under these conditions led to a robust hypoxic response that we measured through TMT-based quantitative mass-spectrometry (Supplementary Table S3) and western blotting (Fig S2). Following 24 hours under hypoxic conditions, proteins involved in glycolysis including SLC2A1, the primary plasma membrane glucose transporter, were significantly upregulated (Fig. S2A). In contrast, subunits of the OXPHOS machinery and the mitochondrial ribosome were significantly decreased, as expected (55). Stabilization of HIF1α was observed within 3 hours (Fig. S2B), as expected. While we found HIGD1A and complex IV subunits were indeed increased in abundance at the early stages of hypoxia (3 hours; Figs. S2B, C), the steady state level of HIGD2A was unaffected (Fig. S2B, compare lanes 1, 2). Finally, we investigated the effects of hypoxia on assembly of complex IV and the supercomplex. Loss of neither HIGD2A nor HIGD1A altered the effects of hypoxia on assembly of the supercomplex (Fig. S2B, lower panels). Moreover, following chronic hypoxia (24 hours) we observed accumulation of the S2 intermediate in both cell lines consistent with the complexes being increasingly sensitive to destabilization in the absence of HIGD2A and HIGD1A. Taken together we conclude that HIGD2A acts on complex IV assembly independent of oxygen levels.

### HIGD2A stabilizes newly translated COX3

Since our results for HIGD1A function were broadly consistent with previous studies, suggesting its involvement in regulating the COX1 module following hypoxia (31), we focused our efforts on clarifying the function of HIGD2A under normoxic conditions. To identify the interaction partners of HIGD2A we used a retroviral system to express FLAG-tagged HIGD2A (HIGD2A^FLAG^) in our HIGD2A^KO^ cell line. Ectopic expression of HIGD2A^FLAG^ led to rescue of supercomplex distribution and an increase in the levels of fully assembled holo complex IV (Fig. 6A). HIGD2A^FLAG^ cells were then solubilized in digitonin and complexes bound to FLAG-affinity gel, following which significantly enriched interactors were identified by affinity enrichment mass-spectrometry (AEMS). As can be seen in Fig. 6B, HIGD2A^FLAG^ associates with subunits of complex IV with mtDNA encoded COX3 (MT-CO3) being the most strongly enriched (Supplementary Table S4). Interestingly, we observed significant enrichment of complex I subunits, subunits of the mitochondrial ribosome (MRPL23, MRPL27 and MRPL50), HIGD1A, complex IV assembly factors COA3 (56–59) and SCO1 (60, 61), and complex I assembly factors NDUFAF2 (39, 62, 63) and NDUFAF4 (64) which is suggestive of the enriched cohort of proteins representing an assembly intermediate rather than the mature complex.

**Figure 6.**
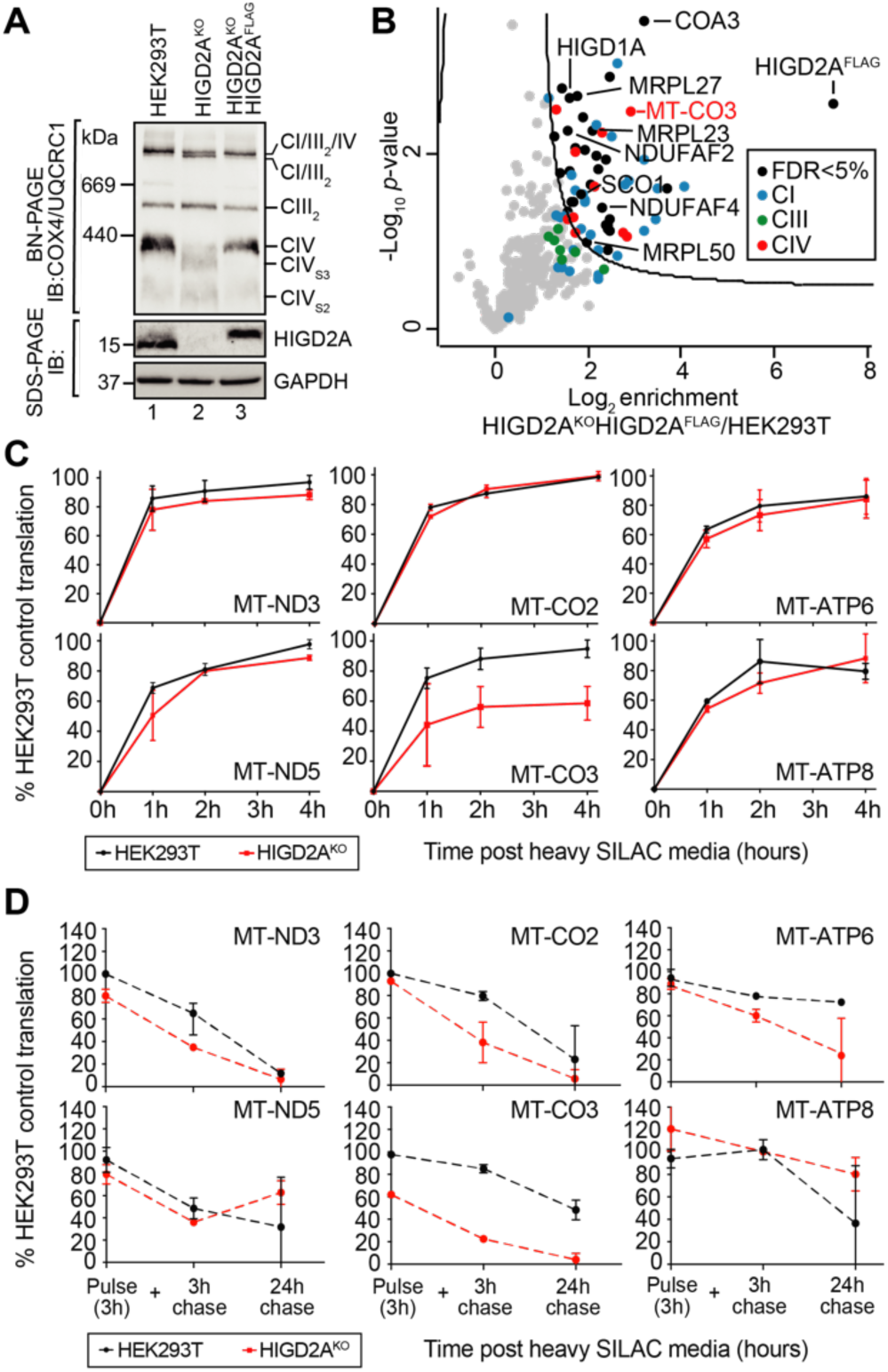
HIGD2A stabilizes newly translated COX3 (A) The HIGD2A^KO^ cell line was complemented with a retroviral HIGD2A^FLAG^ construct and mitochondria analyzed by BN and SDS-PAGE and immunoblotting with the indicated antibodies. (B) Affinity-enrichment mass spectrometry (AE-MS) was performed on the HIGD2A^KO^HIGD2A^FLAG^ cell line and significant interaction partners visualized through a volcano plot. The curved line indicates significance determined through an FDR-based approach (FDR<5%, S0=1). n=3. (C) Pulse SILAC analysis of newly translated mtDNA encoded OXPHOS subunits in HIGD2A^KO^ cells. SILAC media was added following overnight treatment with chloramphenicol (CAP) and analysis was performed at 0, 1, 2- and 4-hours post media addition. Log_2_ transformed heavy peptide-derived intensities were plotted relative to controls. Data reported as mean ± S.D. n=3. (D) As for C but heavy media exchanged with light media after a 3h pulse and cells cultured for the indicated times.

Given the enrichment of the mitochondrial ribosome and the strong association of HIGD2A^FLAG^ with mtDNA-encoded COX3, we asked if mtDNA-encoded subunit translation was affected upon loss of HIGD2A. Typically, translation and stability of mtDNA-encoded subunits is assessed using a radioactive pulse-chase assay and analysis by SDS-PAGE and autoradiography (65). Since COX2 and COX3 have a similar migration on SDS-PAGE (53) we sought to develop a mass-spectrometry based approach capable of clear differentiation between these two proteins. HIGD2A^KO^ and control cells conditioned by overnight treatment with mtDNA translation inhibitor chloramphenicol were pulsed in media containing SILAC (stable isotope labeling by amino acids in cell culture; (66)) amino acids ^13^C_6_^15^N_4_-arginine and ^13^C_6_^15^N_2_-lysine for 1, 2 and 4 hours prior to analysis by targeted mass-spectrometry (described in the methods). Using this technique, we could monitor the translation of 10 out of 13 mtDNA encoded subunits (Supplementary Table S5) including COX2 and COX3. As can be seen in Fig. 6C we observed a significant defect in the stability of newly translated COX3 (MT-CO3) whereas translation of other mtDNA-encoded subunits was comparable to control cells. This result led us to ask if the defect in stability of newly translated COX3 occurs prior to or after membrane insertion. HIGD2A^KO^ and control cells treated as above were pulsed with SILAC media for 3 hours to accumulate newly translated COX3. At the conclusion of the pulse, the amount of newly translated COX3 in HIGD2A^KO^ was ∼60% of that observed in control cells (Figure 6D). The accumulation of other newly translated subunits in HIGD2A^KO^ was similar to control cells, as expected. SILAC media was exchanged with media containing normal amino acids and the stability of SILAC labeled proteins chased for 3 and 24 hours. We observed time dependent turnover of most subunits as is observed in the classic radiolabeled translation assay (67). The turnover rate of newly translated COX3 was similar in both HIGD2A^KO^ and control cells, suggesting the defect in COX3 stability occurs prior to the chase. Moreover, COX3 turnover in HIGD2A^KO^ was similar to the rates observed for other mtDNA encoded proteins observed in both cell lines. Taken together we conclude that the defect in COX3 stability observed in HIGD2A^KO^ occurs early during subunit translation and maturation.

### HIGD2A^KO^ mitochondria contain a supercomplex assembly lacking COX3

Given the apparent retention of the CI/III_2_/IV supercomplex in HIGD2A^KO^ we sought an alternate approach to assess supercomplex content. We utilized a cell line we developed previously that stably expresses the FLAG-tagged complex I subunit NDUFV3 in the corresponding NDUFV3^KO^ cell line (32). NDUFV3 is suggested to be the final subunit added to complex I during its biogenesis and we previously used the resulting NDUFV3^FLAG^ cell line to enrich all detectable subunits of complexes I, III and IV (32) leading us to hypothesize that the cell line can be used to isolate the intact supercomplex. To this end we used CRISPR/Cas9 to generate HIGD2A knockout mutations in the NDUFV3^FLAG^ cell line. HIGD2A^KO^ and NDUFV3^FLAG^HIGD2A^KO^ have comparable defects in complex IV assembly and supercomplex distribution (Fig. 7A). Next, we performed affinity enrichment and mass-spectrometry on both the original NDUFV3^FLAG^ and NDUFV3^FLAG^HIGD2A^KO^ cell lines. While NDUFV3^FLAG^ cells co-isolated complex I, III and IV subunits as expected (Fig. 7B), we also detected HIGD2A, the known complex IV assembly factor COA3 and a number of other mitochondrial housekeeping proteins also detected in our original experiment (Supplementary Table S6 and (32)). We conclude that while the NDUFV3^FLAG^ eluate does not represent the purified supercomplex seen in the recent structures it likely represents a late stage assembly intermediate with a full complement of complex I, III and IV subunits and selected assembly factors. Finally, to assess the impact of HIGD2A loss on the supercomplex we performed the affinity enrichment experiment from NDUFV3^FLAG^HIGD2A^KO^ cells using NDUFV3^FLAG^ as the control cell line. As can be seen in Fig. 7C, while the majority of complex I, III and IV subunits were equally enriched from both cell lines, significantly lower levels of COX3 were enriched from the supercomplex intermediate isolated from cells lacking HIGD2A.

**Figure 7.**
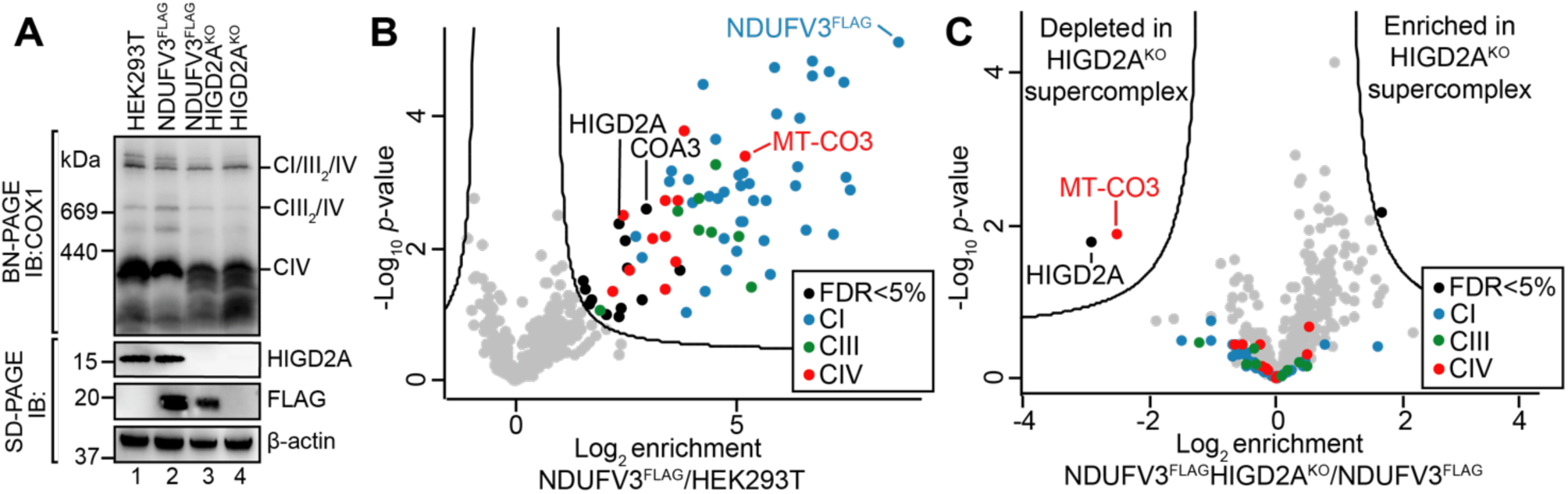
The supercomplex in HIGD2A^KO^ cells lacks COX3. (A) The indicated cell lines were analyzed by BN and SDS-PAGE and immunoblotting with the indicated antibodies. (B) Affinity-enrichment mass spectrometry (AE-MS) was performed on the NDUFV3^FLAG^ and control cell lines and significant FLAG interaction partners visualized through a volcano plot. The curved line indicates significance determined through an FDR-based approach (FDR<5%, S0=1). n=3 independent subcultures. (C) as for B however, the enrichment was performed on NDUFV3^FLAG^/HIGD2A^KO^ relative to the NDUFV3^FLAG^ cell line.

## Discussion

In this study we aimed to clarify the roles of *hypoxia inducible gene 1* domain family members HIGD2A and HIGD1A. Although the mammalian proteins were originally identified in a screen for genes upregulated upon hypoxic stress (29, 30) the majority of work has been focused on their yeast homologs Rcf1 and Rcf2 (named for <u>R</u>espiratory <u>c</u>omplex factors). Three independent groups reported the functions of Rcf1 and Rcf2 as required for the formation of the respiratory chain supercomplexes, which in the yeast *S*. *cerevisiae* are comprised of complexes III and IV (24–26). HIGD2A is thought to be the mammalian orthologue of Rcf1 and knockdown experiments in mouse myoblasts were shown to alter the distribution of complex IV-containing supercomplexes, suggesting the function of HIGD2A in supercomplex assembly is conserved from yeast to man (26). Until now this was the only study to investigate the impact of HIGD2A loss on mammalian complex IV and supercomplex assembly, and as a result the function of HIGD2A has largely been extrapolated from yeast studies. The solution structure of yeast Rcf1 was recently solved by NMR (68) revealing two charged transmembrane helices that form salt bridges leading to formation of a homodimer. Mammalian HIGD2A lacks the charged membrane helices responsible for Rcf1 dimerization, suggesting the functions of Rcf1 and HIGD2A may differ. Despite this point of difference, the results from yeast studies are generally in agreement with our observations. For example, as we found in HIGD2A^KO^ cells, yeast mitochondria lacking Rcf1 still contain the CIII_2_/IV supercomplex (25, 26, 69) (compare to Fig. 3A). Secondly, the association of Cox12 and Cox13 (COX6B and COX6A in mammals) with complex IV is impaired in Rcf1 knockouts as it is in HIGD2A^KO^ cells (Fig. 3B), and likewise the remaining assembly is still capable of association with complex III (24, 25). Finally, there also appears to be a reduction in the steady state levels of COX3 upon deletion of Rcf1 (25, 28, 69). Indeed, in their original study Strogolova and colleagues (25) suggest that Rcf1 may interact with newly translated COX3 to regulate its association with Cox12 (COX6B) and complex IV as a means of modulating complex IV activity. While our data demonstrates a specific defect in the stability of newly translated COX3 (Fig. 6C and D) it remains unclear if HIGD2A regulates translation of the subunit or stabilizes its early stages of biogenesis.

The interaction between HIGD2A, COX3 and other complex IV and respiratory chain subunits detected in our HIGD2A^FLAG^ affinity enrichment experiment (Fig. 6B) argues for HIGD2A being directly involved in adding COX3 to the partly assembled complex, whereas the upregulation of HIGD2A in patient lymphoblasts with an isolated complex IV defect arising from impaired translation of mtDNA-encoded subunits could be taken as an argument for transcriptional regulation (Fig. 2B). Our previous analysis of fibroblasts from a patient with a mutation in a subunit of the mitoribosome and therefore defective translation of all mtDNA-encoded subunits, also showed upregulation of complex IV assembly factors COX15, SCO2, SURF1 (36) suggesting their upregulation is a general response to complex IV dysfunction. Another possibility that warrants investigation is that HIGD2A may work with mitochondrial inner membrane insertase OXA1 (70) to promote membrane insertion of COX3. While OXA1 was not quantified in our HIGD2A^FLAG^ affinity enrichment, we previously showed that OXA1 interacts with HIGD2A as well as several other proteins detected in our experiment (70).

While we cannot completely exclude the possibility that HIGD2A can directly regulate complex IV activity, we did not observe changes in either the levels of HIGD2A or the impact of its loss on complex IV or supercomplex assembly during acute or chronic hypoxia as has been suggested for HIGD1A (Fig. S2). Our conclusion is that the impact of HIGD2A on the supercomplex is a pleotropic effect due to defective complex IV assembly. The intersection of the latter assembly steps of complexes I, III and IV with supercomplex formation (8, 16–18, 20) is also a major factor to be considered in interpretation of the results from ours and other studies. We detected subunits of complexes I, III and IV as well as assembly factors for complexes I and IV in our HIGD2A^FLAG^ affinity enrichment experiment (Fig. 6B), suggesting that HIGD2A acts at the stage of both COX3 module integration into complex IV, and complex IV integration into the supercomplex. Moreover, in our HIGD2A^KO^ cells we observed upregulation of complex I and III subunits found at the interface between complexes I and IV (Fig. 4A). We hypothesize this is a mechanism to compensate for their destabilization following disruption of their interactions with complex IV subunits, in agreement with our previous results demonstrating that loss of the affected complex I subunits leads to total disassembly of the supercomplex and loss of mitochondrial respiration (32). HIGD2A is not the first assembly factor suggested to impact supercomplex assembly indirectly (8). The most well-known example is COX7A2L (SCAFI), which was originally found within the supercomplex but not holo complex IV (6). A role of COX7A2L in supercomplex assembly has been disputed by multiple groups (17–22) and the protein is now understood to act as a checkpoint for assembly of complex III into the supercomplex (17). Another example is UQCC3 (C11orf83), which is an assembly factor suggested to stabilize the CIII_2_/IV form of the supercomplex (71). UQCC3 has also been shown to be involved in early complex III assembly and is found mutated in patients with isolated complex III deficiency (MIM 616097; 72). Importantly, density for neither HIGD2A, COX7A2L nor UQCC3 has been identified in the multiple recent structures representing several different forms of the mammalian supercomplex assembly (1, 2, 4, 7).

As part of this study we also attempted to validate the role of HIGD1A in respiratory chain assembly. The function of mammalian HIGD1A has been studied in more detail than HIGD2A and the protein has been shown to bind subunits of the COX1 module (specifically COX4, COX5A) (31). It has been proposed that during acute hypoxia, the increased amounts of HIGD1A bind complex IV and induce structural changes around the heme *a* active center, increasing complex IV activity (31). In agreement, we also showed upregulation of HIGD1A protein following 3 hours of growth under hypoxic conditions (Fig. S2) as well as upregulation of complex IV subunits, which was previously suggested to be a mechanism of compensation for the reduced oxygen environment (55). Consistent with a general function for HIGD1A in regulation of the respiratory chain during metabolic stress, Ameri and colleagues (73) showed HIGD1A to be upregulated upon glucose deprivation, however under these conditions the authors could only show an interaction with subunits of complex III and not complex IV. In contrast to these studies, under normoxic conditions, HIGD1A was most recently suggested to associate with an early complex IV assembly intermediate representing the nascent COX1 module (10). In agreement with this, we detected HIGD1A along with other complex IV assembly factors and subunits in our HIGD2A^FLAG^ affinity enrichment experiment (Fig. 6B), suggestive of it being associated with complex IV during assembly. However, despite small changes in the levels of COX1 module subunits upon loss of HIGD1A (Fig. 4A, D), the lack of a strong complex IV or supercomplex assembly defect in our HIGD1A^KO^ (Fig. 3A) argues against its role as an assembly factor.

In conclusion, although the hypoxia induced domain family member HIGD2A has previously been implicated in the maintenance of respiratory chain supercomplexes, we find it to be involved in stabilization of mtDNA-encoded COX3. Loss of HIGD2A leads to turnover of nuclear encoded COX3 partner subunits and accumulation of a supercomplex containing a crippled complex IV. In addition to clarifying previous reports suggesting HIGD2A involvement in supercomplex assembly, our study identifies HIGD2A as the first assembly factor required for the biogenesis of the COX3 module of human complex IV.

## Acknowledgments

We thank all members of the Stroud lab for input into experimental design and interpretation of data, Giel van Dooren and Jenni Hayward for advice on flux analysis experiments, the Bio21 Mass Spectrometry and Proteomics Facility (MMSPF) and the Monash Biomedical Proteomics Facility for the provision of instrumentation, training, and technical support. We acknowledge Bice Dibley for assistance in writing scripts used for the annotation and mapping of proteomics data onto Cryo-EM structures. We acknowledge funding from the National Health and Medical Research Council (NHMRC Project Grants 1125390, 1140906 to DAS and MTR and 1068409 to DRT; NHMRC Fellowships 1140851 to DAS and 1155244 to DRT). DHH is supported by a Melbourne International Research Scholarship and Mito Foundation Top-up Scholarship, and HSM by an NHMRC Biomedical Postgraduate Scholarship (1017174). We acknowledge support from the Victorian Government’s Operational Infrastructure Support Program.

## Data Availability

The mass spectrometry proteomics data have been deposited to the ProteomeXchange Consortium via the PRIDE partner repository and the identifier will be made available upon publication.

## Author Contributions

D. Hock and D. Stroud designed the research, D. Hock, B. Reljic, H. Mountford, and A. Compton performed experiments, D. Hock and C-S. Ang developed methodology, all authors analyzed data and wrote the paper.

## Abbreviations

AEMS: affinity enrichment and mass spectrometry
BN-PAGE: Blue Native Polyacrylamide gel electrophoresis
COX: Cytochrome *c* oxidase
ETC: electron transport chain
HIF1α: Hypoxia-inducible factor 1-alpha
HIGD: Hypoxia inducible gene 1
IMM: inner mitochondrial membrane
KO: Knock out of protein expression
MIM: Mendelian Inheritance in Man
mtDNA: mitochondrial DNA
MT-: encoded by mitochondrial DNA
NDUF: NADH dehydrogenase (ubiquinone) oxidoreductase
NMR: Nuclear magnetic resonance
OXPHOS: oxidative phosphorylation
Rcf: Respiratory supercomplex factor
ROS: reactive oxygen species
SCAFI: Supercomplex Assembly Factor I
SILAC: Stable Isotope Labeling with Amino acids in Cell culture
SPS-MS3: Synchronous precursor selection triple stage mass spectrometry
TMT: Tandem Mass Tags
TMPD: N,N,N′,N′-Tetramethyl-p-phenylenediamine

---

**Supplementary Figure S1.**
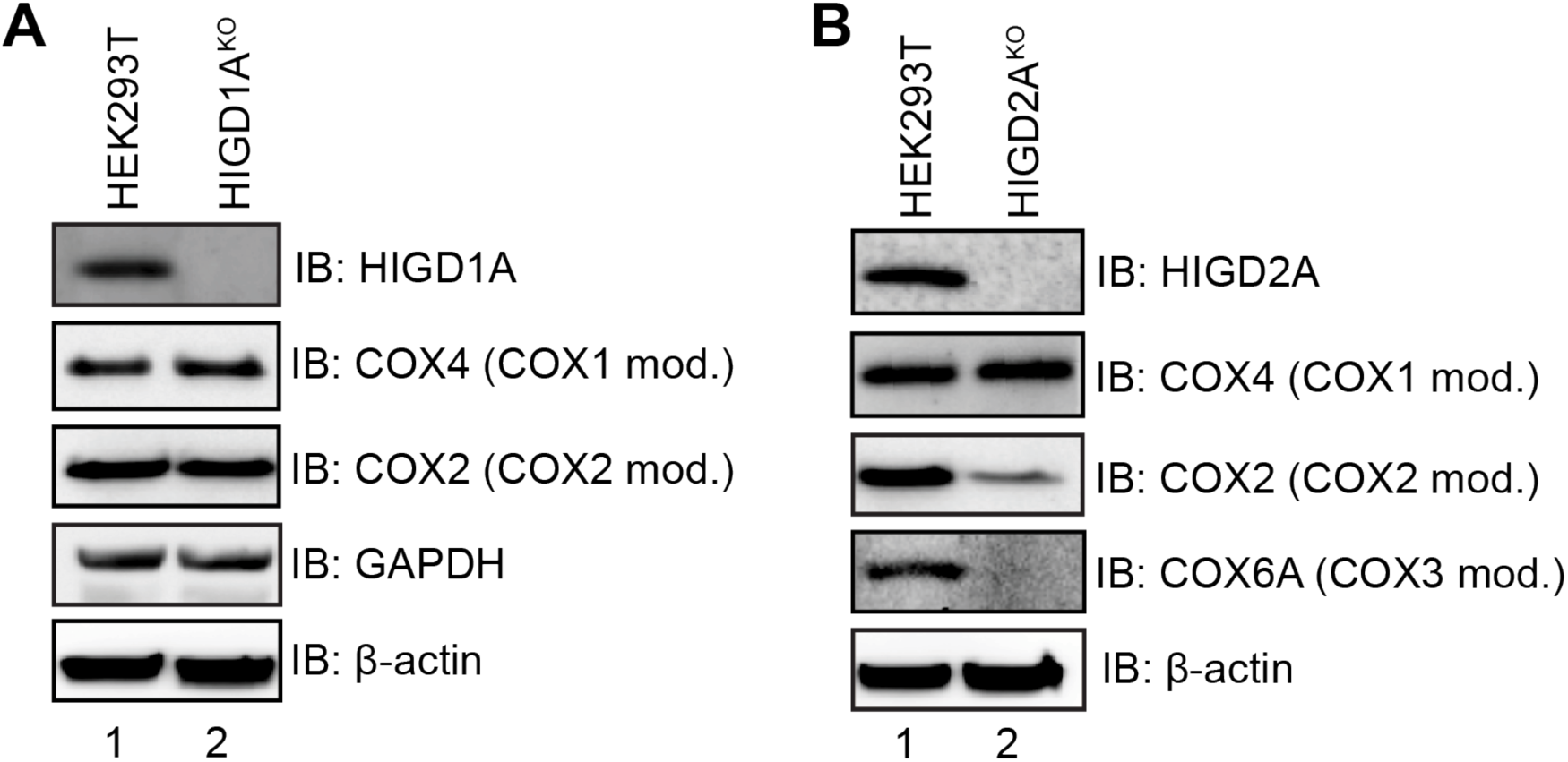
Loss of HIGD2A leads to reduced levels of COX2 and COX6A. (A) HIGD1A^KO^ cells were analyzed by SDS-PAGE and immunoblotting with the indicated antibodies. (B) HIGD2A^KO^ cells were analyzed by SDS-PAGE and immunoblotting with the indicated antibodies. Mod., complex IV module.

**Supplementary Figure S2.**
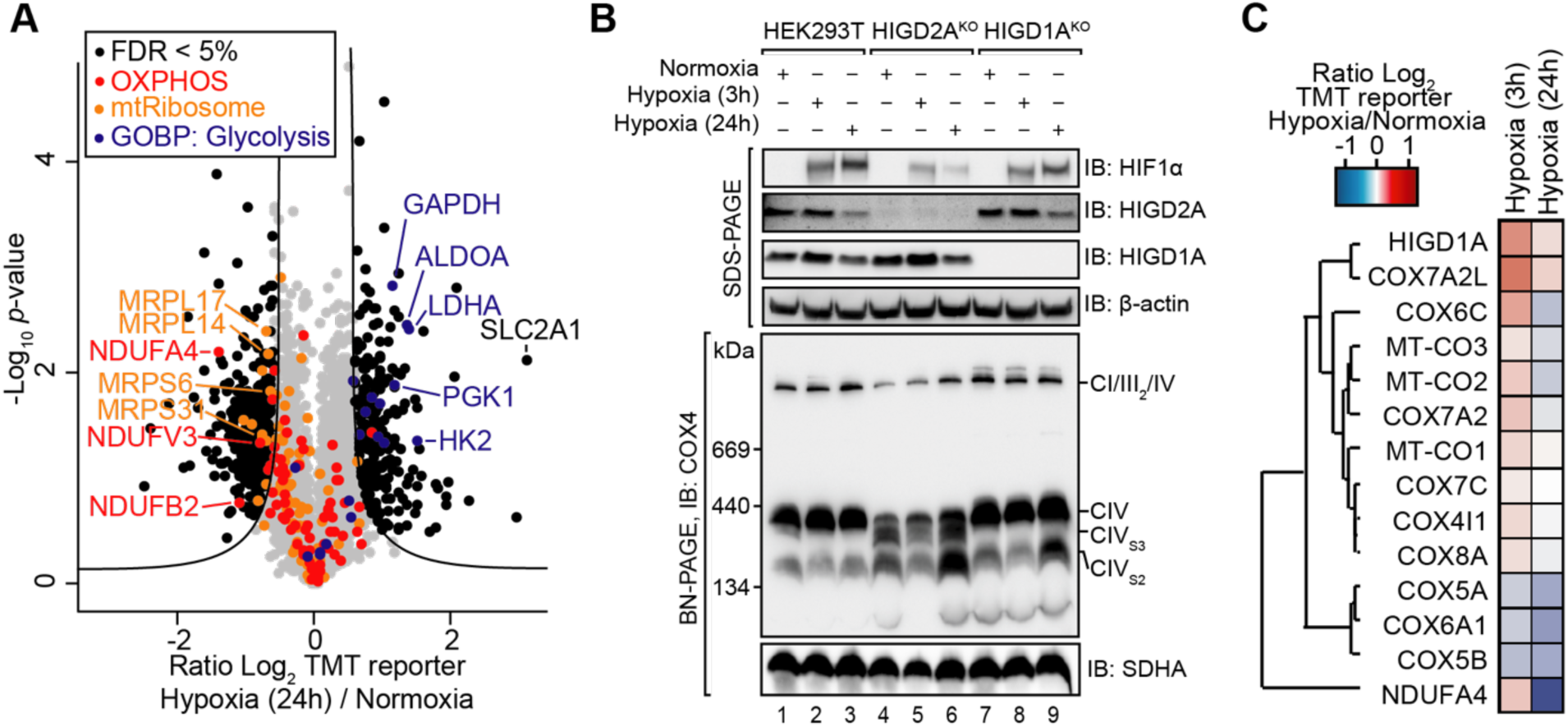
Hypoxia induces HIGD1A but not HIGD2A. (A) Volcano plots depicting relative levels of cellular proteins in HEK293T cells following 24 h of growth under hypoxic conditions. Black dots and the curved line represent significantly altered protein abundance determined through a false discovery rate-based approach (5% FDR, s0=1). N=2 independent subcultures. GOBP, proteins contained within the Gene Ontology: Biological Process category. (B) Cells were grown under normoxia and the hypoxic conditions for the indicated times. Cell lysates were analyzed by SDS- and BN-PAGE and immunoblotting with the indicated antibodies. (C) The relative levels of complex IV subunits in HEK293T cells following 3 or 24 h of growth in hypoxia were subjected to hierarchical clustering and presented as a heatmap.

